# Deep in the Bowel: Highly Interpretable Neural Encoder-Decoder Networks Predict Gut Metabolites from Gut Microbiome

**DOI:** 10.1101/686394

**Authors:** Vuong Le, Thomas P. Quinn, Truyen Tran, Svetha Venkatesh

**Affiliations:** Applied AI Institute, Deakin University, Geelong, Australia

## Abstract

Technological advances in next-generation sequencing (NGS) and chromatographic assays [e.g., liquid chromatography mass spectrometry (LC-MS)] have made it possible to identify thousands of microbe and metabolite species, and to measure their relative abundance. In this paper, we propose a sparse neural encoder-decoder network to predict metabolite abundances from microbe abundances. Using paired data from a cohort of inflammatory bowel disease (IBD) patients, we show that our neural encoder-decoder model outperforms linear univariate and multivariate methods in terms of accuracy, sparsity, and stability. Importantly, we show that our neural encoder-decoder model is not simply a black box designed to maximize predictive accuracy. Rather, the network’s hidden layer (i.e., the latent space, comprised only of sparsely weighted microbe counts) actually captures key microbe-metabolite relationships that are themselves clinically meaningful. Although this hidden layer is learned without any knowledge of the patient’s diagnosis, we show that the learned latent features are structured in a way that predicts IBD and treatment status with high accuracy. By imposing a non-negative weights constraint, the network becomes a directed graph where each downstream node is interpretable as the additive combination of the upstream nodes. Here, the middle layer comprises distinct microbe-metabolite axes that relate key microbial biomarkers with metabolite biomarkers. By pre-processing the microbiome and metabolome data using compositional data analysis methods, we ensure that our proposed multi-omics workflow will generalize to any pair of -omics data. To the best of our knowledge, this work is the first application of neural encoder-decoders for the interpretable integration of multi-omics biological data.

## 1 Introduction

The human gut is a complex ecosystem in which host cells and foreign organisms coexist, cooperate, and compete. Suspended in this ecosystem is a milieu of nutrient metabolites that act like a currency, being exchanged and converted by the organisms living in the environment. Technological advances in next-generation sequencing (NGS) and chromatographic assays [e.g., liquid chromatography mass spectrometry (LC-MS)] have made it possible to identify thousands of microbe and metabolite species, and to measure their relative abundance. By applying NGS and LC-MS on fecal samples, one gains two complementary “views” on the complex ecosystem in which gut bacteria produce, consume, and induce the metabolic milieu. These data modalities have each advanced our understanding of elusive gut pathologies like inflammatory bowel disease [37], and are increasingly being collected in parallel [40, 22, 43].

Once rarely encountered, inflammatory bowel disease (IBD) has become a major health burden in developed countries, with its incidence steadily rising since the second world war [25]. IBD is an umbrella term for two distinct clinical syndromes, Crohn’s disease (CD) and ulcerative colitis (UC), that are both characterised by a chronic immunological disturbance in the gastrointestinal (GI) tract, caused by genetic and environmental factors [14, 15]. While CD presents with patchy transmural (deep) inflammation in any part of the GI tract, UC is marked by diffuse mucosal (superficial) inflammation that extends from the rectum through the colon [42]. Although the inflammation in IBD does not have an infectious origin, patients with CD and UC exhibit an irregular gut microbiome, having less bacterial diversity, a depletion of healthy bacteria, and an excess of unhealthy bacteria [25, 7, 12]. These changes have been partly attributed to an abnormal immune response to benign commensual organisms [25]. Microbiome irregularity, called dysbiosis, also associates with concurrent changes in microbial functions [28] and in metabolic profiles [19], that together can disrupt normal gut physiology. For example, Marchesi et al. found low levels of short chain fatty acids in IBD fecal samples [26], which may be related to changes in how the gut bacteria metabolize carbohydrates [37].

Franzosa et al. studied the paired microbial and metabolic profiles of 164 IBD patients and 56 healthy controls, producing one of the largest publicly available multi-omics data set of its kind [10]. In their multi-omics analysis, the authors report that only 6% of all possible pairwise associations were statistically significant, concluding that metabolites “tend not to associate mechanistically” with the microbiome [10]. However, multi-omics data integration can be approached in several ways, ranging from simple to complex. We organize these approaches into four tiers. The first, and simplest, uses iterative univariate-univariate regressions, e.g., measuring the Pearson’s correlation between a single bacteria and a single metabolite (as done by Franzosa et al. [10]). This straightforward method is implemented in several microbiome-specific software tools [44]. Although pair-wise associations can be easy to interpret, they lack the ability to model the additive effect of bacteria (or metabolite) co-occurence. The second uses iterative univariate-multivariate regressions, e.g., measuring a single bacteria as a function of all metabolites (and *vice versa*). This method is still easy to interpret, and has been applied to infer gene expression from DNA mutations [11], as well as metabolite variables from the microbiome [40]. The third uses a single multivariate-multivariate regression, such as a canonical correlation (CanCor) analysis. CanCor is a powerful tool that can find a combination of microbes that maximally correlate with a combination of metabolites. This technique has been applied previously to study the relationship between volatile breath metabolites and gut microbiome in IBD patients [38], but its widespread application is limited by high-dimensionality [37] (at least without regularization [27]).

Stepping further toward deeper modeling, we explore the fourth tier which leverages a multi-layer neural network to model a single multivariate-multivariate regression. The hidden layers of these networks act like a switchboard to connect the input layer with the output layer through a set of intermediate nodes that can learn complex (non-linear) patterns between the layers. In biology, deep neural networks have been used in many applications, including predicting the expression of 20,000 genes from 1,000 hallmark genes [4]. While useful, the hidden layers of these networks do not have a natural meaning that connects the model to real biological processes. To address this limitation, many neural networks follow the *encoder-decoder* pattern in which the network has an *hourglass* shape, featuring a narrow middle layer that compresses the input-output relationship. This layer is regularized to be low dimensional so that it only has enough information space to describe the input-output transformation. As such, all other information is filtered out. This layer divides the network into two specialized parts: the **encoder** and the **decoder**. One could think of a generic encoder-decoder network as a neural “signal translator”, designed to turn one data set **X** into another data set **Y**, through a middle representation **Z**.

Encoder-decoder networks have been studied in computer vision in the form of fully convolutional models [23], where an image is encoded into a compact represetation that is then decoded into the desired feature map. In biomedical imaging, U-net [35] has succeeded in segmenting cell images with the encoder and decoder forming the two interacting shafts of a U shape. In biology, autoencoders – a special type of encoder-decoder architecture where the input and output are identical – have been used to cluster yeast, Pseudomonas, and cancer, where the hidden layer supposedly provides a biologically meaningful abstraction of the data [5]. However, generic encoder-decoders that transform one data domain to another have not apparently been used for multi-omics data integration. In this generalized form, encoder-decoders can act like a CanCor, predicting multiple outputs from multiple inputs through a latent space. Unlike CanCor, encoder-decoders learn deep non-linear relationships between the features.

Using the encoder-decoder architecture, we seek to find a good model that can provide simple, descriptive, and verifiable patterns within the multi-omics data. In this manuscript, we introduce our highly interpretable neural encoder-decoder design that can learn the non-linear and synergistic relationship between the gut microbiome and their surrounding metabolites. In doing so, we demonstrate that (a) neural networks outperform linear models in microbiome-metabolome predictions, and that (b) network sparsification, along with a non-negative weights constraint, improves the accuracy, stability, and interpretability of the encoder-decoder model. Importantly, we show that our neural encoder-decoder model is not simply a black box designed to maximize predictive accuracy. Rather, the network’s hidden layer (i.e., the latent space, comprised only of sparsely weighted microbe counts) actually captures key microbe-metabolite relationships that are themselves clinically meaningful. Although this hidden layer is learned without any knowledge of the patient’s diagnosis, we show that the learned latent features are structured in a way that predicts IBD and treatment status with high accuracy. Taken together, our work demonstrates that paired multi-omics data can be integrated using a neural network whose hidden layer abstracts a clinically meaningful representation of the input data without any supervision. By pre-processing the data using standard compositional methods, we ensure that our encoder-decoder workflow will generalize to any pair of -omics data.

## 2 Methods

### 2.1 Data acquisition and processing

We acquired paired microbiome and metabolome data as raw proportions from the supplement of Franzosa et al. [10]. To reduce the dimensionality of the data, we removed features in which more than 50% of the measurements were zero. We replaced the remaining zeros with a very small number using the `cmultRepl` zero-replacement function implemented in `zCompositions`, an imputation tool which explicitly models the relative nature of NGS and LC-MS data [30]. Next, we processed the data using one two of pipelines: “Complete” or “Summarized”.

In the “Complete” pipeline, we applied the centered log-ratio (clr) transformation directly to the bacteria species-level and metabolite cluster-level abundances. The clr is a cornerstone in the analysis of compositional data [1, 2, 8, 32]:

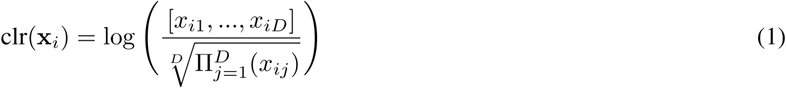

where x_*i*_ is a sample vector of bacteria or metabolite abundances. Beyond transforming the data into real numbers, the clr is also convenient for machine learning applications because the “normalization” factor is applied to each sample independent of all other samples, thus preserving test set independence.

In the “Summarized” pipeline, we aggregated the bacteria species-level abundance into genus-level abundance by summing across the respective genus members. We also aggregated the metabolite cluster-level abundance into class-level abundance by summing across the respective class members. We retrieved the species-to-genus conversion table from the Integrated Taxonomic Information System (ITIS) (via the the R package `taxize` [3]), and the metabolite-to-class conversion table from the supplement of Franzosa et al. [10]. Features that did not belong to any genus or class were dropped. Table 1 describes the dimensionality of the data before and after processing.

**Table 1:**
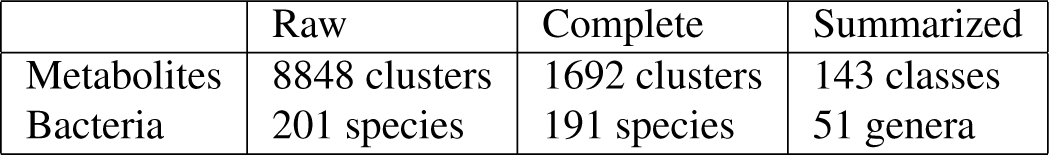
This table counts the number of features in the data before and after processing. Removal of features with mostly zeros reduced the number of metabolite clusters from 8848 to 1692, and the number of bacteria species from 201 to 191. Summarization further reduced the dimensionality to 143 classes and 51 genera.

|  | Raw | Complete | Summarized |
| --- | --- | --- | --- |
| Metabolites | 8848 clusters | 1692 clusters | 143 classes |
| Bacteria | 201 species | 191 species | 51 genera |

### 2.2 Our Motivation: Predicting Metabolites from Microbes

The nature of the relationship between the two data modalities can be explicitly represented if we can find a way to predict one from the other. The ideal model would not only identify correlations between the pairs of data, but would also reveal the mechanism through which they influence each other. These complex processes can be considered from the point-of-view of regression models that can predict metabolite abundance from bacteria abundance.

We formulate the prediction problem as a search for the parametric function *f*, with parameter set *θ*, that takes as input the (clr-transformed) microbe abundances *X* to estimate the (clr-transformed) predicted metabolite abundances *Ŷ* of the real (clr-transformed) observations *Y* :

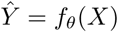

where 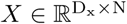 contains *D*_*x*_ microbes and 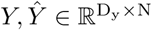 contains *D*_*y*_ metabolites for the same *N* subjects.

The parameter set *θ* is estimated by minimizing the error between the predicted and real observations, e.g., using the mean square error (MSE):

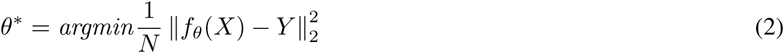

The choice of the function *f* and the optimization strategy used to solve Equation 2 is the key to having a predictive model with better accuracy, stability and interpretability. In the next section, we will propose a deep neural network that is specifically designed for this goal. First, lest us consider the most direct and straightforward baseline to model the functional *f* : a linear regression (LR) model, where the abundance of each metabolite is predicted as a linear combination of all available microbes:

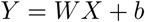

where 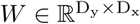 is the linear transformation matrix and *b* is the bias term. The problem in Equation 2 now has the form:

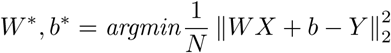

While being the simplest model, LR suffers from operating on a full transformation matrix. The large number of parameters (i.e., weights) not only makes the LR prone to overfit when the data set is small and high-dimensional, but also makes the model difficult to interpret. To reduce the density of the linear transformation matrix, Lasso *L*_0_*/L*_1_ regularization constraints can be added to reduce the number of active (i.e., non-zero) weights in the transformation matrix *W* [41]. With this regularization, the objective function turns into

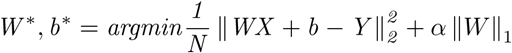

where the hyperparameter *α* controls the sparsity of the model and ‖*W* ‖_1_ is *l*_1_-norm of the linear weights. An example Lasso model is illustrated in Figure 1b.

**Figure 1:**
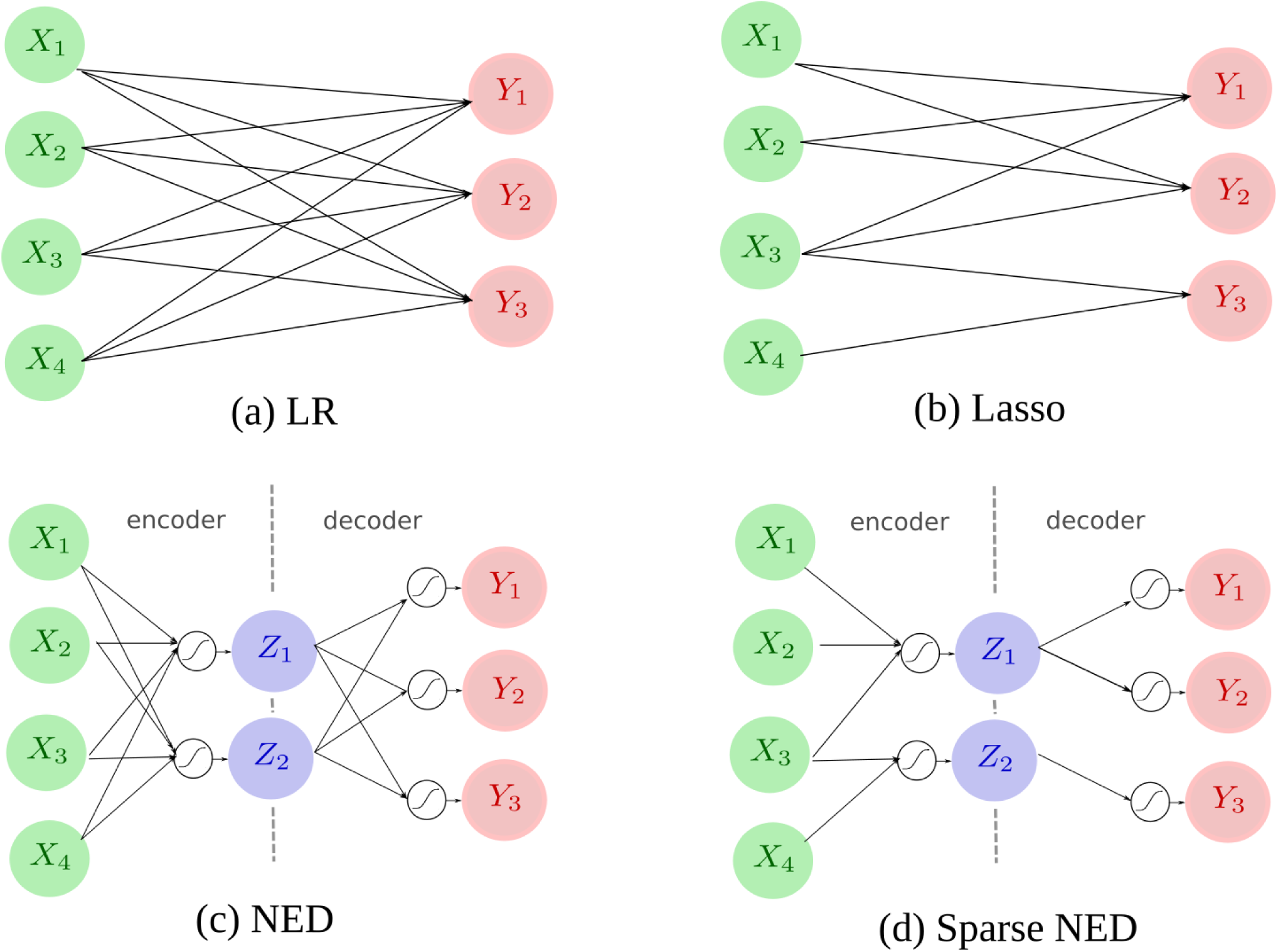
This figure compares the computation network of NEDs with their direct linear counterparts. The green circles are components of metabolite abundances {*X*_*i*_}, the red circles are components of bacteria abundance {*Y*_*j*_}, and the blue circles are latent variables {*Z*_*k*_}. Arrows denote linear combinations from the source to the destination, while white circles with the tanh graph denote non-linear activation steps.

### 2.3 Our Model: A Sparse Neural Encoder-Decoder for Data Integration

Although simple and easy to interprete, a univariate multi-omics model depends on the major assumption that the processes in which microbes affect metabolites are singular and independent from each other. On the other hand, neural networks can learn many sub-processes of a global process that governs the dynamic, multi-stage interaction between the two modalities.

The traditional method for learning multivariate-multivariate relationships is cannonical correlation (CanCor) analysis, though like LR it can only find linear relationships between the data modalities. Instead, we propose to construct a robust and interpretable deep neural network. Our model is designed to relax the key assumption behind LR and CCA: that there exists a direct and linear relationship between the two data modalities. This relaxation will extend our representation of the predictive model via two hypotheses:

1. There exists intermediate factors that act in the middle of the process that transforms microbes to metabolites.
2. The transformation between these factors may contain non-linear parts.

#### 2.3.1 The Neural Encoder-Decoder (NED) Network

The neural encoder-decoder (NED) architecture aims to predict a multivariate random process using the information from another multivariate process through an intermediate representation called the **latent feature space**. The part of the network that extracts relevant information from the input (i.e., microbes) into the latent space is called the **encoder**. The part of the network that predicts the output (i.e., metabolites) from the latent space is called the **decoder**. The latent feature space is realized as the narrow **hidden layer** lying between the encoder and decoder. Compared with a direct linear model like linear regression, the encoder-decoder network should have a more robust representation because the weights undergo non-linear activation. As such, we expect the NED to perform better.

To maintain the interpretability of the model, and to make it more robust to small amounts of training data, we restrict the number of hidden layers to one. Our initial experiments show that a large number of hidden layers does not improve the predictive performance of the NED because the model is easily overfitted on the limited training data.

In operation, the two parts of the model work together sequentially. First, the **encoder** extracts the relevant information from the microbes *X* to store in the latent variables *Z* by an encoding function consists of a linear kernel *W*_*e*_ followed by a fixed non-linear activation function *σ*_*e*_:

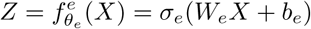

Second, the **decoder** decrypts the latent content in *Z* to predict the value of *Y* as *Ŷ* using a similar decoding function:

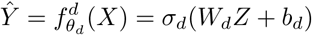

The final predictive model is the composition of the encoder and decoder:

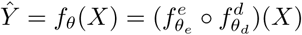

This model is trained using the loss function:

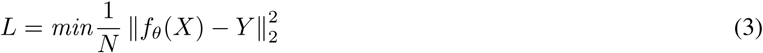

with the optimized parameter set *θ*:

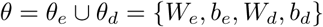

Figure 1 compares the computation network of NEDs with their direct linear counterparts. Let 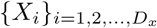 denote the *D*_*x*_ components of *X*, and let 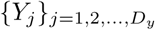 denote the *D*_*y*_ components of *Y*. The number of latent variables *D*_*z*_ is a meta-parameter chosen by heuristics. To increase robustness and interpretability, models with fewer connections are preferred. An LR model has *C*_LR_ = *D*_*x*_.*D*_*y*_ connections, while an NED has *C*_NED_ = *D*_*x*_.*D*_*z*_ + *D*_*z*_.*D*_*y*_ connections. Thus, we see that when *D*_*z*_ << *D*_*x*_, *D*_*y*_, then *C*_NED_ < *C*_LR_. This makes NED less likely to overfit and easier to interpret. We can further reduce the density of NED networks using a sparsity procedure that eliminates redundant connections between the layers.

#### 2.3.2 Sparsifying the NED Network

To further improve the stability and interpretability of the model, we seek to learn an NED model with the fewest number of active weights: a **sparser neural network** where most weights equal zero. Research into sparser neural networks have achieved major advancements recently. This line of work is motivated by the intuition that deep networks are generally over-complete, and sparse networks could reduce computational overhead [9]. Furthermore, sparser deep networks are also known to ease the explanability of the underlying process [39, 33]. The major approaches to sparser networks work by either pruning unnecessary weights [13] or enforcing sparsity contraints as an additional regulatory loss [24]. To sparsify NED into a new model, called **Sparse-NED**, we use a pre-training screening method that is similar to the one-shot pruning strategy first prosposed by Lee et al. in [20].

The learning of Sparse-NED consists of two stages: screening and training. In the *screening stage*, we identify the connections that are most useful in extracting the information needed to predict metabolite abundance from microbe abundance. All other links are marked as unnecessary. In the *training stage*, the network is trained with these redundant links deactivated from the forward and backward operations.

The screening stage starts with one training iteration on the fully connected model, where the derivative *g*(*w*; *D*) of the loss *L* (from Equation 3) is estimated through back-propagation on a sample set of data *D*. These derivatives are the key to evaluating the importance of a connection because when the derivative at a connection has a high magnitude, that connection will have a measurable effect on the loss, and hence be more salient to the prediction of metabolite abundance. Slightly different from [20] – where only a mini-batch is used as *D* for sensitivity calculation – we use all of the available training data instead. The saliency of each connection *s*_*c*_ is calulated as the normalized magnitude of the derivatives:

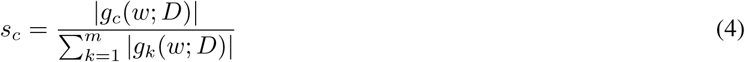

Next, the connections in the network are sorted in descending order of *s*_*c*_, and only top-*k* connections are kept for network training and inference. The number of kept connections *k* is controlled through the sparsity level *β* relative to total number of connections in the fully connected network.

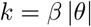

The hyperparameter *β* is to balance the accuracy and sparsity of the model. In our experiment, we choose a *β* so that the number of active (i.e., non-zero) connections is on par with other sparse models in the comparison. Using the remaining connections, the Sparse-NED is trained via back-propagation. An example Sparse-NED is illustrated in Figure 1d.

#### 2.3.3 The Non-Negative Weights Constraint

With a small number of active connections, Sparse-NED is significantly easier to understand than a fully connected NED. However, among the remaining connections, many of them spontaneously have negative weights. These negative connections in neural networks have been understood to inhitbit an analysis of how the factors from one layer affect another [6]. To improve the interpretability of a network, non-negative contraints can be used to prevent the weights from falling below zero [45]. In our application, the non-negative weights provide a clearer meaning to how microbe abundances contribute to metabolite abundances, both in encoder and decoder, because the contribution is always positive.

To implement this insight, we apply the non-negative contrainst on Sparse-NED model by clamping the intermediate estimation of the parameters to [0, *∞*) at every training iteration:

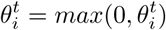

where 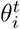 is the *i*^*th*^parameter of the model at iteration *t* in training.

In our experiment, we observe that non-negative constraints not only promote interpretability, but also have a beneficial effect on the overall fitness and sparsity of the model. When weights are forced to be positive or zero, the nodes in each layer tend to compete with each other for influence. This actually makes the Sparse-NED *even more sparse*.

### 2.4 Model Evaluation

With the goal of deeply understanding the bacteria-metabolite relationship, we want a model that is not only accurate, but also highly interpretable, and whose interpretation is stable across different folds of the data. In this section, we discuss the three criteria used to evaluate our predictive models: accuracy, sparsity, and stability.

#### Accuracy

The accuracy of a predictive model *f*_*θ*_ is calculated by the Pearson correlation coefficient between the predicted metabolite abundance *Ŷ* = *f*_*θ*_ (*X*) and the measured abundance *Y* for the top 10 best predicted metabolites. Compared to intensity-based criteria, the correlation coefficient is not influenced by the scales used in the signal and is reliable across different data normalization methods. However, to mitigate the influence of spurious correlation, correlations are always calculated using the clr-transformed data.

#### Sparsity

The sparsity of *f*_*θ*_ is measured by the number of active linear connections 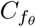 in the model. In the case of NED and its variations, it includes the weights from both the encoder and decoder network sections.

#### Stability

While accuracy and sparsity are desired properties of a predictive model, their performance and interpretation are only reliable when they are consistent across changes in the training set composition. For each model family, we evaluate the stability of predictive models by measuring the average pair-wise similarity [16] of the model parameters *θ* across different training sets. Specifically, we divide the data set into 5 folds and learn an instance of the model for each f old. T hen, for each pair of model instances, we calculate the Pearson’s correlation coefficient between the model parameters. The main measurement for the stability of a method, the *stability index*, is calculated as the average similarity between all pairs of model instances within the model family.

To concentrate on the stability of the model architecture, we calculate the stability index using the binary adjacency matrix of the connections between each layer in the model. For fully connected models (e.g., a linear regression or CanCor) – where every factor in a layer affects every factor in the subsequent layer – the binary adjacency matrix contains all ones. Therefore, the stability index always equals one and is not worth mentioning. For sparse methods (e.g., Lasso and NED), the stability index measures the consistency and reliability of the cross-layer connections. For multi-layer models like NED and its variants, we use the first model instance to initialize the training of the other instances so that we preserve the correspondence of the mid-layer variables (e.g., the latent variable “V5” in fold 1 is the same as the latent variable “V5” in fold 2).

### 2.5 Interpretation of Network Layers

The neural network contains three layers: the microbe input layer, the hidden (i.e., latent) layer, and the metabolite output layer. In order to understand the nature of the latent space, we performed a separate analysis of each layer and compared their results. All microbe and metabolite analyses were performed on the clr-transformed data, while the latent space analyses were performed on the unaltered (i.e., tanh-compressed) data.

#### 2.5.1 Differential Abundance (DA) Analysis

We performed an FDR-adjusted analysis of variance (**ANOVA**) of the three experimental groups [ulcerative colitis (**UC**), Crohn’s disease (**CD**), and healthy control (**HC**)] for the microbe, metabolite, and latent space feature sets (separately). We consider any feature with a p-value *p* < 0.05 to be significant. Note that the prior clr-transformation makes univariate statistical testing of relative data valid, so long as the results get interpretted with regard to the reference used [8, 32].

#### 2.5.2 Redundancy Analysis (RDA)

We performed a redundancy analysis (**RDA**) of the microbe, metabolite, and latent space feature sets (separately) using the `rda` function from the `vegan` R package [29].

#### 2.5.3 Random Forests

We used the microbe, metabolite, and latenet space features to train a random forest classifier to predict several two-factor outcomes (see Results and Discussion). Random forest models were trained using the `randomForest` function from the `randomForest` R package [21] without any feature selection or hyper-parameter turning. For each feature space, and for each outcome, we compared the average “out-of-the-box” AUC across 25 randomly sub-sampled test set splits using the `plMonteCarlo` function from the `exprso` R package [31].

## 3 Results and Discussion

### 3.1 Summarization preserves data structure

The data sets produced by high-throughput molecular assays like NGS and LC-MS often have many more features than samples. In statistics, high-dimensionality is a problem because the likelihood of a false discovery increases with each additional test. In machine learning, high-dimensionality increases the likelihood of an overfit and also makes the resultant model more difficult to interpret. Therefore, it is sensibile to reduce the number of features before training a model. After removing zero-laden features, we used domain-knowledge to aggregate the remaining features into biologically meaningful groups. For metabolites, we used functional classes; for bacteria, we used assigned genus. Table 1 describes the dimensionality of the data before and after processing.

Figure 2 shows the first two principal components for the metabolite data (top) and microbiome data (bottom), as processed using the “Complete” (left) and “Summarized” (right) pipelines. Here, we see that using domain-knowledge to summarize the metabolite clusters into classes, and the bacteria species into genera, does not appear to alter the fundamental structure of the data. Otherwise, we note that the IBD patients cluster more distinctly accoridng to their gut metabolites than their gut bacteria. Indeed, a differential abundance analysis of the metabolic data reveals 128 (out of 143) significant metabolite classes, compared with 15 (out of 51) significant bacterial genera (see Supplement for a complete list). Table 2 shows the average clr-transformed abundances for select bacteria genera, chosen because they were found previously to associate with IBD. Two genera, **Ruminococcus** and **Fusobacterium**, show an association that agrees with past literature [18].

**Table 2:** This table shows the average clr-transformed abundances for select bacteria genera, chosen because they were found previously to associate with IBD. Since abundance is expressed relative to the per-sample mean, positive values signify above-average presence for samples within that group. The evidence for previous association is taken from [18]. All p-values are FDR-adjusted. Two genera, **Ruminococcus** and **Fusobacterium**, show an association that agrees with past literature.

| Genus | p-value | Control | IBD | UC | CD | Previous Association | Agreement |
| --- | --- | --- | --- | --- | --- | --- | --- |
| Clostridium | <0.001 | 3.785 | 4.898 | 4.397 | 5.331 | Decreased in IBD | No |
| Ruminococcus | <0.001 | 2.895 | -0.352 | 0.517 | -1.102 | Decreased in IBD | Strong |
| Lactobacillus | 0.006 | 1.352 | 2.202 | 1.586 | 2.733 | Decreased in IBD | No |
| Fusobacterium | 0.012 | -2.56 | -1.866 | -2.336 | -1.46 | Present in IBD | Strong |
| Bifidobacterium | 0.097 | 3.179 | 3.417 | 4.262 | 2.686 | Decreased in IBD | Mixed |
| Bacteroides | 0.199 | 5.698 | 4.939 | 5.142 | 4.764 | Decreased in IBD | Weak |

**Figure 2:**
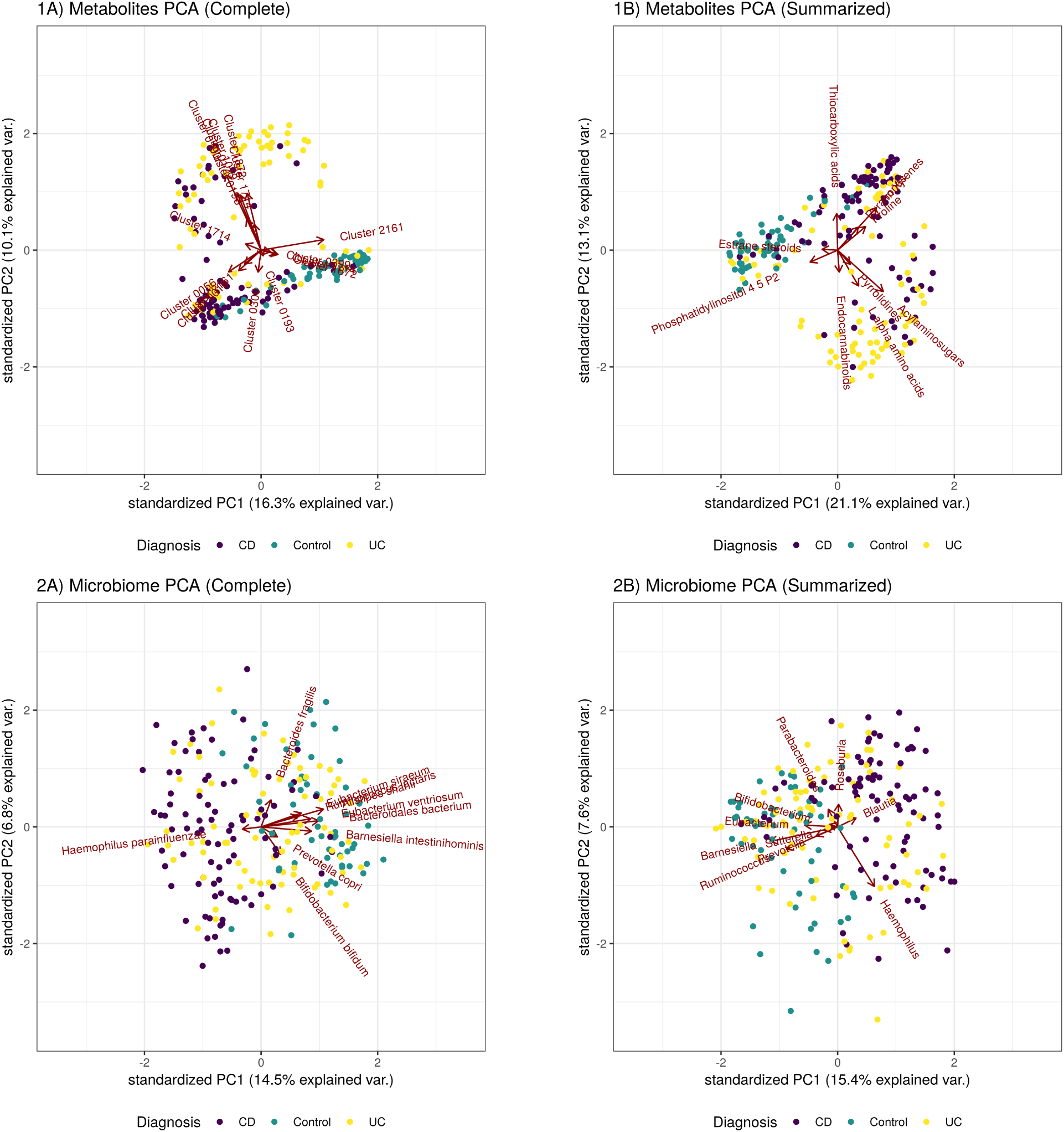
This figure shows the first two principal components for the metabolite data (top) and microbiome data (bottom), as processed using the “Complete” (left) and “Summarized” (right) pipelines. Here, we see that using domain-knowledge to summarize the metabolite clusters into classes, and the bacteria species into genera, does not appear to alter the fundamental structure of the data.

### 3.2 Microbe abundance predicts metabolite abundance

Since metabolites and bacteria both associate with IBD, it is meaningful to explore their mutual dependence. In the Introduction, we described four approaches to integrating multi-omics data: univariate-univariate regression, univariate-multivariate regression, multivariate-multivariate regression, and neural networks. Franzosa et al. reported weak univariate-univariate regressions for these data [10]. Here, we evaluate the performance of the other multi-omics approaches by benchmarking the performance of microbe-metabolite predictive models with 5-fold cross-validation. The quantitative evaluation is done using the three criteria described in Section 2.4.

#### 3.2.1 Neural Encoder-Decoders outperform linear regression

We do not expect that the microbiome alone can predict the abundance of all metabolites. Rather, we are motivated to answer two research questions: (1) Which metabolites can be predicted by the microbiome? and (2) How reliable is that prediction? Table 3 and Table 4 show the performance of the microbe-metabolite prediction models for the “Complete” and “Summarized” data, respectively. The tables are organized by the multi-omics integration scheme used: univariate-multivariate, multivariate-multivariate, or neural network. To answer the two research questions, we compare the accuracy of each model for the top 10 best predictions, along with its sparsity and stability.

**Table 3:** Performance when predicting metabolite (cluster-level) abundance from microbiome (species-level) abundance. Acronyms: NED neural encoder-decoder; E encoder; D decoder. The stability indices of the fully connected models are naturally 1.0 and not worth including.

| *Method* | *Top 10 Corr.Coeff* | *Active links* | *Stability index* |
| --- | --- | --- | --- |
| Linear Regression (LR) | 0.46 | 241956 | — |
| + Sparsity (LASSO) | 0.70 | 6143 | 0.50 |
| + Non-neg. weights | 0.70 | 4417* | 0.52 |
| CCA | 0.47 | 20449 (E) + 241956 (D) | — |
| Linear NED | 0.60 | 10010 (E) + 118440 (D) | — |
| + Nonlinear activation (NED) | 0.74 | 10010 (E) + 118440 (D) | — |
| + Sparsity (Sparse-NED) | 0.74 | 2216 (E) + 4284 (D) | **0.79** |
| + Non-neg. weights (Nonneg-Sparse-NED) | **0.74** | **1522\* (E) + 2568\* (D)** | 0.63 |

**Table 4:** Performance when predicting metabolite (class-level) abundance from microbiome (genus-level) abundance. Acronyms: NED neural encoder-decoder; E encoder; D decoder. The stability indices of the fully connected models are naturally 1.0 and not worth including.

| *Method* | *Top 10 Acc Corr.Coeff* | *Active links* | *Stability index* |
| --- | --- | --- | --- |
| Linear Regression (LR) | 0.39 | 9741 | — |
| + Sparsity (LASSO) | 0.62 | 898 | 0.57 |
| + Non-neg. weights | 0.58 | 685 | 0.61 |
| CCA | 0.61 | 2601 (E) + 9741 (D) | — |
| Linear NED | 0.59 | 7150 (E) + 84600 (D) | — |
| + Nonlinear activation (NED) | 0.63 | 7150 (E) + 84600 (D) | — |
| + Sparsity (Sparse-NED) | **0.67** | 551 (E) + 702 (D) | **0.80** |
| + Non-neg. weights (Nonneg-Sparse-NED) | 0.66 | **360\* (E) + 428\* (D)** | 0.66 |

#### 3.2.2 Sparsity improves accuracy and interpretability

Without regularization, it is easy for a model to overfit. Therefore, it is no suprise that the sparse linear regression (i.e., LASSO) and the sparse neural encoder-decoder outperform their fully connected counterparts. Nevertheless, in both cases, the neural encoder-decoder outperforms linear regression (although LASSO performs impressively well).

#### 3.2.3 Non-negative weights improve interpretability

With the default linear regression and the neural encoder-decoder network, weights can take on any value. Here, we propose using a non-negative weights constraint to improve the interpretability of the model. For a neural network, this constraint means that when an input node contributes to a hidden node, that contribution is always additive. This allows us to interpret the hidden layer as an *aggregation* of the input. Likewise, each output node is computed by an aggregation of hidden nodes.

Combined with sparsity, the non-negative weights constraint ensures that each node is the simple sum of just a few elements. Table 3 and Table 4 show that this constraint does not seriously reduce accuracy, though it advantageously does force the network to be *even more sparse* (see *Active links* column).

### 3.3 The latent space is clinically coherent

The latent space of the encoder-deconoder network is a learned abstraction of the input data, designed to describe how gut microbes associate with gut metabolites for all patients. As such, variance within the hidden layer reflects inter-patient variance within the microbe-metabolite axis. We focus this section on the **sparse and non-negative neural encoder-decoder model**, trained using the “Summarized” data, because we think it nicely balances accuracy with interpretability. For this neural network, the value of each hidden node equals tanh(x · w + *b*), where **x** is the per-sample microbe abundances, **w** is the weights associated with each microbe, and *b* is an offset. Together, these hidden nodes comprise a new feature space, learned in a fully unsupervised way, that can be analyzed directly with routine statistical modelling.

#### 3.3.1 The latent space associates with IBD

For each of the 70 nodes in the hidden layer, we can compute its variance across all patients: some nodes are more variable than other nodes. We can also compute the amount of weight coming in and out of each node: some nodes have more “traffic” than other nodes. High-traffic nodes strongly relate multiple microbes to multiple metabolites, while low-traffic nodes describe fewer or subtler relationships. Figure 3 plots the node variance against the node traffic. Here, we see that the high-variance nodes are usually the high-traffic nodes, suggesting that the nodes which model the most microbe-metabolite interactions also vary the most between individual patients. An ANOVA of the latent features reveals that the high-variance-high-traffic nodes significantly associate with IBD (FDR-adjusted p-value < 0.05). Since the latent space is an abstraction of the relationship between microbes and metabolites, its association with IBD suggests that the relationship between microbes and metabolites is itself associated with IBD.

**Figure 3:**
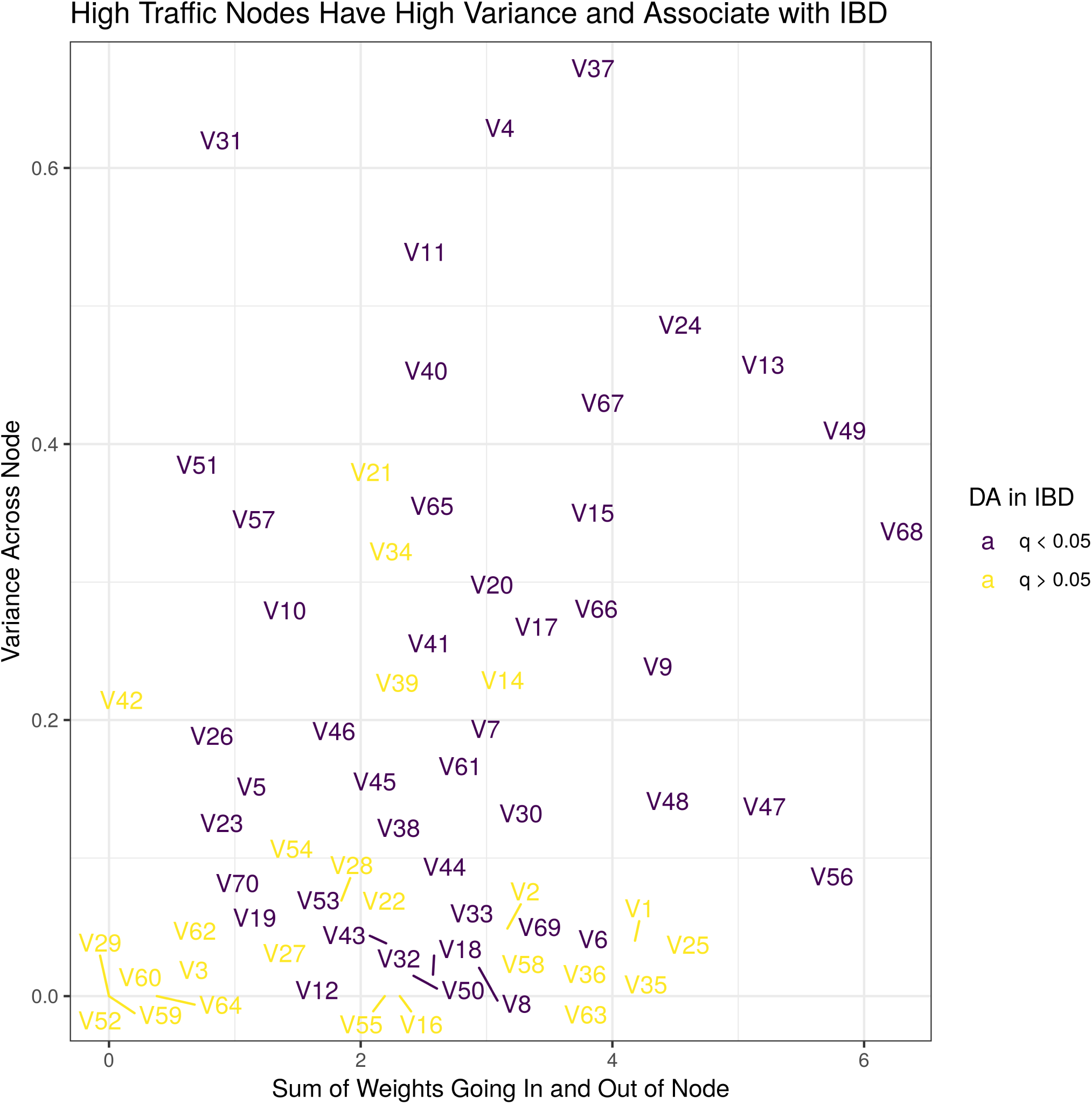
This figure shows the variance of a hidden layer node (y-axis) versus the total weight going in and out of that node (x-axis). There are 70 nodes, named arbitrarily, and colored by their association with IBD (FDR-adjusted p-value < 0.05). The most heavily weighted nodes are the most important for predicting metabolite abundances. The most variable nodes differ the most between patients. Here, we see that the most variable and most heavily weighted nodes all associate with IBD.

#### 3.3.2 The latent space is a noise filter

While an ANOVA allows us to measure the statistical association between the latent space and IBD, we can further understand the clinical relevance of the latent space with a redundancy analysis (RDA). RDA is a principal component analysis that constrains the principal axes so that they describe the part of the latent space that is also explained a “conditioning matrix”, **L**. Here, the conditioning matrix contains the clinical covariates: **age, fecal calprotectin, diagnosis, antibiotic use, immunosupressant use, mesalamine use**, and **steroid use**. In the left panel of Figure 4, we see that the first RDA axis involves multiple correlated nodes that all associate strongly with the IBD diagnosis (CD vs. HC vs. UC). Although medication use is confounded by diagnosis, we see that the off-axis node “V21” is perfectly anti-correlated with antibiotic use (and CD), while the off-axis node “V66” is correlated with it. The right panel shows box plots for some of the longest first-axis and off-axis arrows. By looking at the RDA eigenvalues, we know that 26.9% of the latent space can be explained by the clinical covariates in **L**. This is more than the 12% of the microbe data, and the 23.6% of the metabolite data, explained by **L**. In other words, a higher fraction of the hidden layer is explained by clinical covariates than the input or output layer. This finding supports two hypotheses. First, the relationship between microbes and metabolites is itself associated with IBD. Second, the encoder-decoder latent space acts like a noise filter that can extract the clinically relevant part of the source data in a fully unsupervised way.

**Figure 4:**
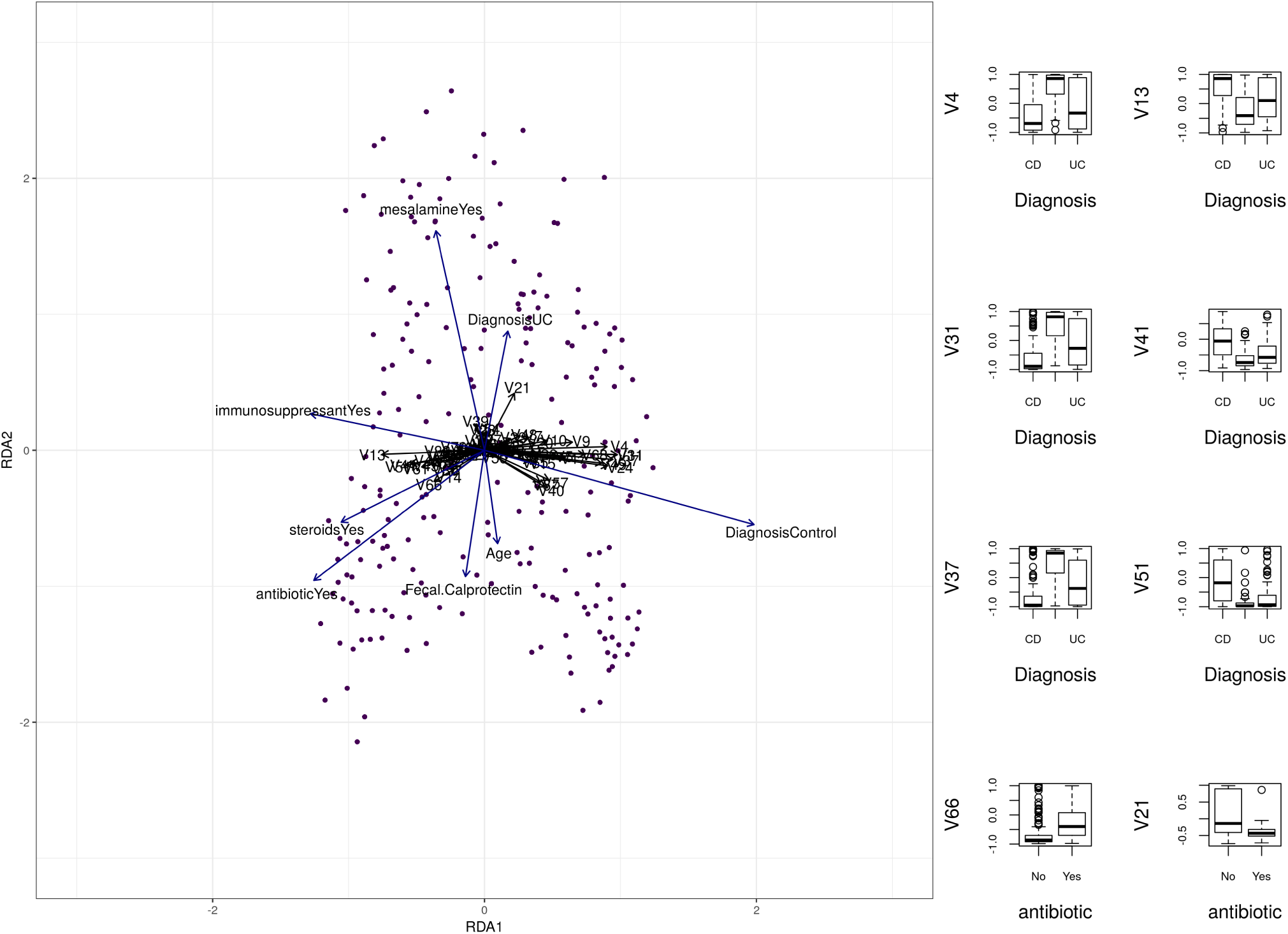
The left panel shows the first two RDA axes of the latent feature space, constrained by the known clinical covariates: age, fecal calprotectin, diagnosis, antibiotic use, immunosupressant use, mesalamine use, and steroid use. The right panel shows box plots for some of the longest first-axis and off-axis arrows. Several hidden nodes describe clinical events.

#### 3.3.3 The latent space is discriminative

From our differential abundance analyses, we know that the microbes, metabolites, and latent features all associate with IBD (though there are more metabolite associations than microbe associations). Although we have shown that the latent space is clinically coherent, we want to further demonstrate its discriminative power in classification tasks. Table 5 shows the average “out-of-the-box” AUC for binary classifiers trained on 25 randomly sub-sampled training sets. For most outcomes, the latent space classifier performs at least as well as the microbe classifier. However, when predicting antibiotic use and immunosuppressant use, the hidden layer is actually *more predictive* than either the microbe and metabolite abundances.

**Table 5:**
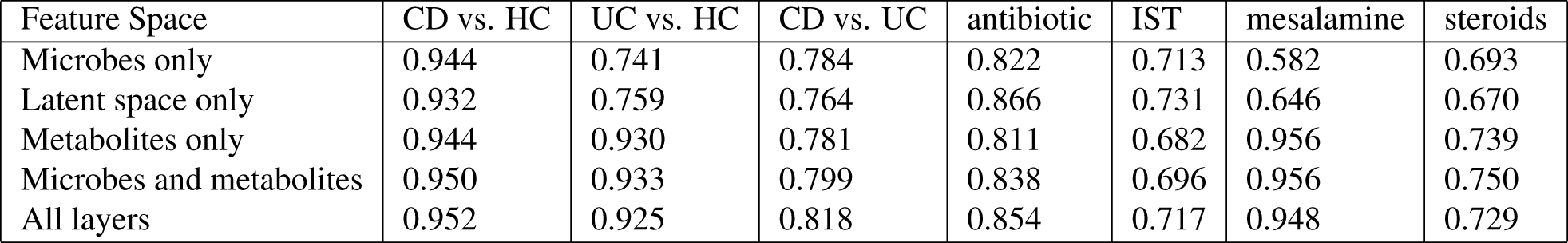
This table shows the average “out-of-the-box” AUC for binary classifiers trained on 25 randomly sub-sampled training sets. In most cases, the latent space classifier performs at least as well as the microbe classifier. Acronyms: CD Crohn’s disease; UC ulcerative colitis; HC healthy control; IST immunosuppressive therapy.

#### 3.3.4 The latent space is interpretable

Figure 5 shows a three layer graph relating microbes to metabolites, built using the edge overlap across all 5 training set folds. The middle layer contains latent variables that weigh the microbe abundances so that they maximally predict the metabolite abundances. The graph reveals a general structure: the top half describes how the microbes that are enriched in healthy guts predict the metabolites that are also enriched in healthy guts, while the bottom half describes how the microbes that are depleted in healthy guts predict the metabolites that are also depleted in healthy guts. **Ruminococcus** and **Fusobacterium** are both replicated IBD biomarkers, and the latent variables that relate them to metabolites also associate with IBD.

**Figure 5:**
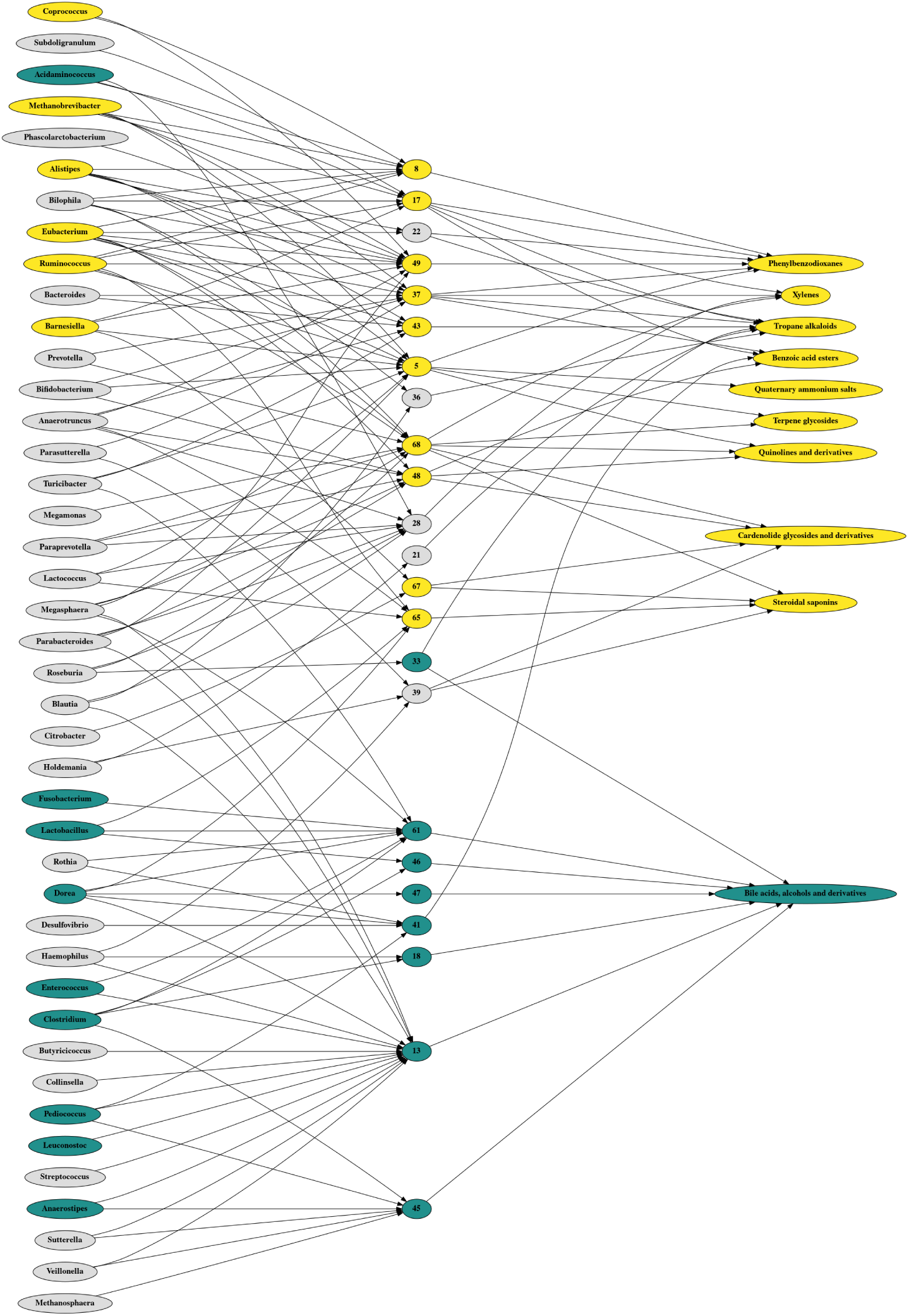
This figure shows a three layer graph relating microbe abundances to the top 10 best predicted metabolite abundances, built using the edge overlap across all 5 model instances. Yellow nodes are significantly enriched in healthy guts, while blue nodes are significantly enriched in IBD guts. The middle layer contains latent variables that weigh the bacteria abundances so that they maximally predict the metabolite abundances. Most of these latent variables, learned in a fully unsupervised way, are themselves significantly associated with IBD.

In the upper graph, we see how **Ruminococcus** (among others) contributes to 6 latent variables which go on to predict several healthy metabolite signatures, including **tropane alkaloids** and **steroidal saponins**, as well as other plant-derived compounds. It is interesting, though perhaps not surprising, that some of the plant-derived compounds enriched in the healthy gut have known medicinal properties [17, 36]. In the lower graph, we see how **Fusobacterium** (among others) contributes to a single latent variable; this node, V61, is highly predictive of the abundance of **bile acids, alcohols, and deriatives**. This finding is consistent with the literature which suggests that the bile acid conjugate taurine is a substrate for bacteria metabolism, and that a defect in the detoxification of taurine by-products is associated with ulcerative colitis [34].

## 4 Summary

Inflammatory bowel disease (IBD) presents a major health burden to developed countries. Although IBD is not infectious, patients with Crohn’s disease (CD) and ulcerative colitis (UC) exhibit an abnormal gut microbiome as well as an altered gut metabolome. In this manuscript, we propose a neural encoder-decoder model to learn a set of weighted connections that can predict metabolite abundances using only microbe abundances. We show that this neural network outperforms linear models for microbiome-metabolome predictions, and that sparsification, along with a non-negative weights constraint, further improves the accuracy, stability, and interpretability of the encoder-decoder model. Importantly, the neural encoder-decoder model is not simply a black box designed to maximize predictive accuracy. Rather, the hidden layer of the model can help visualize the predictive relationship between microbes and metabolites. Moreover, the learned latent feature space (i.e., the hidden nodes themselves) appears to structure the data in a clinically coherent way: the latent space associates with, and predicts, IBD diagnosis and medication use. Our finding suggests that the microbe-metabolite axis itself, not just the microbes and metabolites alone, is an IBD-specific biomarker signature. To the best of our knowledge, this work is the first application of neural encoder-decoders for the interpretable integration of multi-omics biological data.

## 5 Declarations

### 5.1 Ethics approval and consent to participate

Not applicable.

### 5.2 Availability of data and material

The neural encoder-decoder model is available from https://github.com/vuongle2/BiomeNED. The transformed data used to train the model, the latent feature space, and the differential abundance results are available from https://doi.org/10.5281/zenodo.3255151.

### 5.5 Authors’ contributions

VL and TPQ conceptualized the project and drafted the manuscript. VL implemented the neural encoder-decoder model and performed the benchmark analyses. TPQ reviewed the IBD literature, prepared the data, and performed the analysis of the latent space. TT helped design the neural network. TT and SV supervised the project. All authors revised and approved the final version of the manuscript.

## 5.6 Acknowledgements

Not applicable.

